# Decline in *Kappaphycus alvarezii* invasion in the Gulf of Mannar, India

**DOI:** 10.64898/2026.08.04.742698

**Authors:** Yamini Srikanth, Sandeep Pulla, Naveen Namboothri, Elrika D’Souza

## Abstract

Blue Economy models position aquaculture as a key pathway to securing global food security. Species selected for aquaculture typically show rapid growth, high stress tolerance and fast biomass accumulation, but these same traits may increase their potential to become invasive when introduced beyond their native range. We investigated the invasion history and current status of the commercially important red seaweed *Kappaphycus alvarezii* in the Palk Bay–Gulf of Mannar region of India. This is one of the world’s largest cultivation hubs, a climatically vulnerable marine biodiversity hotspot, and one of the three regions to report invasion. We combined in-water surveys, interviews with wild seaweed collectors, and a review of published literature to reconstruct the history of invasion and assess current status. Invasion has declined substantially, with interviews indicating that the disappearance of invasive populations began around 2014. We discuss several non-mutually exclusive explanations for this decline, including climate change, loss of coral substrate, herbivory, and reduced vitality of the seaweed. Although the decline in invasion is encouraging for coral reefs, our findings raise questions about the ecological and socioeconomic consequences of introducing non-native aquaculture species under Blue Economy initiatives, particularly in ecologically sensitive regions vulnerable to climate change.

## 1. Introduction

Oceans are the new frontier for development, and the Blue Economy has emerged as a global framework for promoting sustainable economic growth, creating livelihoods, enhancing food security, and supporting climate change mitigation through the use of ocean resources (Bennett et al. 2019). This includes sectors like fisheries, renewable energy, and increasingly, aquaculture of finfish and macroalgae. Although various initiatives that can be considered part of the Blue Economy have been implemented for decades, the framework has recently gained considerable traction. Multilateral initiatives like the United Nations Sustainable Development Goals, particularly SDG 14 (Life Below Water), and agreements and frameworks advanced by the UN Decade of Ocean Science for Sustainable Development have supported this momentum (UNRIC 2022). Blue Economy schemes are increasingly promoted across Asia, Africa, and Small Island Developing States (Bennett et al. 2019), regions which are particularly vulnerable to the effects of climate change (Gill et al. 2023).

Macroalgae have become central to the Blue Economy, even being considered the “keystone” due to their varied applications in food, feed, biofuels, and carbon sequestration (Cai et al. 2021; Ismail and Zokm 2025). However, the aquaculture sector poses a significant risk to native biodiversity and ecosystem functioning as the industry relies on translocating and cultivating Non-Native Species (NNS) (Namboothri et al. 2012; Oficialdegui et al. 2025). Examples of marine bioinvasions with severe ecological impacts include the Indo-Pacific lionfish, brought to the Western Atlantic and Caribbean through the aquarium trade (Côté and Smith 2018), and comb jellies, introduced to the Mediterranean by the release of ballast water (Jaspers et al. 2018). Seaweeds such as *Codium fragile*, and *Caulerpa recemosa var. cylindrica* have also had significant impacts (Chapman et al. 2006; Neill et al. 2006). Species introduced for aquaculture are of particular concern as the traits that make them good aquaculture species also increase their risk of bioinvasion, namely fast growth, short reproductive cycles, and stress tolerance (Namboothri et al. 2012; Oficialdegui et al. 2025).

One marine alga that has formed a critical component of a number of Blue Economy schemes is *Kappaphycus alvarezii*, a red alga that is prized for its ability to grow extremely quickly, doubling or even tripling its biomass in a period of 45-60 days (Hayashi et al. 2017). *K. alvarezii* produces the industrially significant thickener carrageenan, which is used in a wide variety of consumer goods from food products to cosmetics (Hayashi et al. 2017). The total value of the seaweed products industry is more than 6 billion USD (Fatima Ferdouse et al. 2018), and a number of national and international schemes have been implemented to promote the widespread cultivation of *K. alvarezii* (Hayashi et al. 2017). *K. alvarezii* is a non-native species (NNS) in most cultivation regions. However, marine biological invasions of NNS has caused significant ecological damage, driving biodiversity loss, ecosystem change, and altering species interactions (Oficialdegui et al. 2025).

*Kappaphycus alvarezii* is native to the Philippines, where its cultivation began in the early 1970s. Its rapid growth and the utility of carrageenan in various industrial products led to its introduction to various tropical and subtropical regions globally (Hayashi et al. 2017). In 1983, it was first reported as invasive in Hawaii, having escaped from cultivation rafts (Russell 1983). By 2005, it had been listed as “one of the world’s 100 worst invasives” (Invasive Species Specialist Group 2005). Another invasion was reported from Panama in 2013 (Sellers et al. 2014), and in our study region of the Gulf of Mannar, India in 2005 (Pereira and Velecar 2005).

The Gulf of Mannar lies along the southeast coast of India, close to Sri Lanka, with 21 small islands surrounded by fringing reefs. These reefs are a part of the Gulf of Mannar Marine National Park (GMMNP), established to conserve the significant biodiversity of the area. The region is heavily affected by a number of stressors, the most significant of which is climate change, with warming-induced bleaching causing sharp declines in coral cover (Raj et al. 2021). Storms and cyclones have also shaped the region, with 2 significant cyclonic storms since 2002 (IMD, 2020). Approximately 41,000 fisher families depend on this marine landscape for livelihoods, with an estimated number of 12,981 traditional, small-scale fishers (Robert Panipilla and Marirajan T 2014). There is a high degree of natural resource dependence, and a need to support communities in a time of increasing uncertainty due to climate change.

*Kappaphycus alvarezii* was introduced to this region for cultivation in 2002, and was subsequently reported as invasive (Arasamuthu A. et al. 2023). Researchers highlighted its ability to overgrow and “smother” corals (Kamalakannan et al. 2014). The status of the invasion has been reported sporadically, with limited primary ecological data. Due to the absence of long-term monitoring data from the Gulf of Mannar, India, the contemporary status of invasion in the region was not known.

As an important aquaculture hub and one of three invaded regions globally, examining the current status of *K. alvarezii* invasion in India is vital to understand both dynamics of invasion and to inform future cultivation strategies.We drew from published literature, interviews with wild seaweed collectors who free dive in the region, and primary ecological surveys to examine the current status of invasion and to place a timeline to the trajectory of invasion in the Gulf of Mannar.

## 2. Methods

### 2.1. Study region

The Gulf of Mannar and the Palk Bay lie along the south-east coast of India (Panel 2, Figure 1), in close proximity to Sri Lanka. Cultivation of *K. alvarezii* began in the Palk Bay in 2002 (Panel 2, Figure 1), and existing literature on *K. alvarezii* invasion in this region has largely documented invasion within the Gulf of Mannar Marine National Park (GMMNP). The Gulf of Mannar and Palk Bay are connected by the Pamban Pass, and the movement of *K. alvarezii* from the cultivation region of the Palk Bay into the Gulf of Mannar is thought to be the source of invasion. The GMMNP is managed and administered by the Tamil Nadu Forest Department (TNFD) under the Ministry of Environment, Forest and Climate Change, Government of India. The park consists of 21 islands, stretching from Rameswaram in the East to Tuticorin in the West. These islands have been divided into four island groups, and these groups are separated by distances of between 10–50 kilometers. Several of these islands are surrounded by fringing coral reefs, although the structure and composition of the reefs differ based on the bathymetry of the islands.

**Fig. 1.**
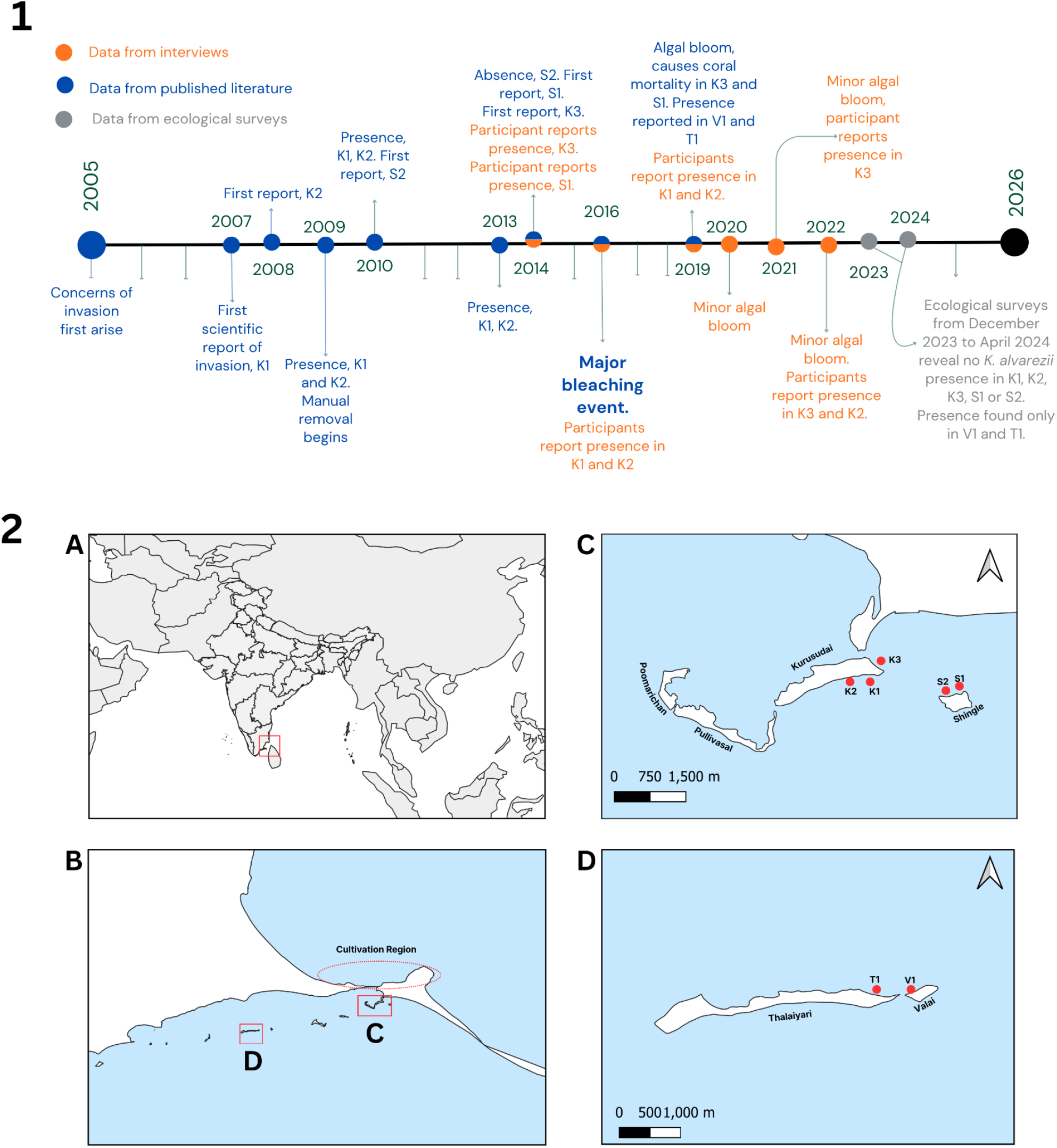
Panel 1: Timeline depicting key events in the history of invasion of *K. alvarezii*. Timeline consolidates information from three sources: interviews, published literature and ecological field surveys Panel 2: A: Study region within Asia B: Sampling sites C and D within the Gulf of Mannar, and the cultivation region in Palk Bay C: Sampling Sites in the first island group D: Sampling sites in the third island group

### 2.1. Literature Review

We performed a literature review to collate, compare, and identify emergent themes in invasion literature. In order to understand dynamics and patterns of *K. alvarezii* invasion, we reviewed published literature from the three areas it has been reported as invasive: Kāne’ohe Bay (Hawaii), Bocas del Toro (Panama), and the Gulf of Mannar (India). This information was collected exclusively from published scientific literature. Due to the paucity of peer-reviewed sources from the Gulf of Mannar, not all published scientific literature included in this review was peer-reviewed. Literature was collated from Google Scholar, and the full list of search terms can be found in the supplementary (Supplementary 1). All papers from 1970 (when *K. alvarezii* cultivation began) to the present were included in the review, though only a selection of key references are provided here. A complete list of references can be found in the supplementary (Supplementary 2).

### 2.2 Semi-Structured Interviews

Interviews with seaweed collectors were conducted in order to contextualise the often-contradicting narrative found within published literature. Interviews were semi-structured, and open-ended. Seaweed collectors are a community that have been diving and collecting wild seaweed around the Gulf of Mannar. Different seaweeds are harvested for a variety of purposes from fertilizers to dyes. Collectors harvest them based on purpose and sale price. They were the target group of our surveys for a few key reasons. Unlike fishers who typically remain on fishing boats, seaweed collectors free dive on reefs and interact with reefs visually. Based on preliminary interactions, we determined that members of this community could likely identify most algal species on sight. Questions were posed to participants and responses were open-ended (the complete interview in English and Tamil is accessible in Supplementary 2). In addition to questions, visual cues of fish and coral species, as well as maps, were utilised to better validate survey responses. Participants were only interviewed regarding presence in the first island group, as the second, third and fourth island groups are not commonly visited by seaweed collectors.

We combined snowball sampling and key informant sampling methods. We first identified and interviewed key informants who are well-known as seaweed collectors. These informants recommended additional participants. This approach allowed us to start with well-informed participants and progressively reach a broader sample gaining both depth from key informants and breadth from the expanding network of participants. A broader sample size also helped us triangulate the information generated through key-informant interviews. All interviews were conducted between February to May 2024.

Human ethics clearance (EC Number-NCF-EC-05/01/2024-(89)) was obtained from the Nature Conservation Foundation, Mysuru, India. Informed prior consent was obtained from participants. An interview form, with information regarding anonymity of responses, and that interviews would be recorded and transcribed, was composed in Tamil (ref. Supplementary 3). This was verbally read out to the participants. The interviews were conducted only with the explicit consent of the participants.

### 2.3 Ecological Field Surveys

During the first iteration of the literature review, we identified a number of invaded sites in the Gulf of Mannar, where *K. alvarezii* had been reported to colonize reefs. These invaded sites and proximate reefs were a priority during ecological surveys.

To quantify the extent of *K. alvarezii* invasion, we conducted in-water visual surveys to confirm *K. alvarezii* presence or absence. These visual surveys covered the first island group. We estimate we covered upwards of 75% of all accessible reefs (safe diving sites with no strong currents), recording presence/absence in all sites reported as invasive within this island group, and all nearby reefs.

To quantify the cover of *K. alvarezii*, we selected 6 sites within the GMMNP. Site selection was based on the in-water visual surveys, and conversations with the TNFD. The TNFD informed us of two sites in the third island group with small invaded patches, separated by a distance of 500 meters. We measured benthic composition and *K. alvarezii* cover using photo-quadrats. We placed 3 quadrats of a 50×50 cm size at fixed intervals along three, 10 m transects. The length was set at 10 m due to the relatively small size of the invaded sites. This led to a total of nine quadrats per site. We quantified the percentage of area covered by different benthic categories: sand, live coral, branching live coral, dead coral, branching dead coral, coral rubble, crustose coralline algae, other macroalgae, and *K. alvarezii*, by overlaying 50 points onto each photo-quadrat using ImageJ. These six sites were divided into two categories based on the findings of this survey. Formerly invaded sites were those that had reported *K. alvarezii* in the past, but where it was not observed during our surveys. Currently invaded sites were those that presently have *K. alvarezii*.

## 3. Results

### 3.1 Insights from the literature review

The literature review revealed a detailed timeline of *K. alvarezii* invasion

#### 3.1.1 Initial documentation and establishment (2005-2008)

*K. alvarezii* cultivation was introduced to the Palk Bay-Gulf of Mannar region in 2002 (Panel 1, Figure 1). Three years after this introduction, Pereira and Verlecar (2005) reported *K. alvarezii* growing in shallow subtidal regions outside of cultivation. In 2008, the first comprehensive study established that it had gained extensive cover in Kurusudai Island, within the first island group (Chandrasekaran et al. 2008; Balasubramanian et al. 2014). It was also reported from the two neighboring islands of Shingle and Poomarichan (Edward and Bhatt 2012). The exact distribution of the algae was unreported at the time, but it appeared to occupy different patches of different reefs (Balasubramanian et al. 2014). The invasion was of significant concern to the Ministry of Environment, Forest and Climate Change. Consequently, a ban on *K. alvarezii* cultivation was enforced in 2008, prohibiting the cultivation of *K. alvarezii* within the Gulf of Mannar region, though it was still permitted in the Palk Bay (Mandal et al. 2010; Krishnan et al. 2021).

#### 3.1.2 Management interventions and ecological observations (2008-2019)

No new invaded sites were reported during this period. Nonetheless, management interventions in the form of manual removal were attempted. Free divers would remove accessible *K. alvarezii* from reefs, and all the collected seaweed would be transported off site. Between 2012 and 2014, repeated cycles of manual removal were attempted by the TNFD (Kamalakannan et al. 2014). It was concluded that manual removal alone was insufficient to control *K. alvarezii* spread (Kamalakannan et al. 2014).

#### 3.1.3 New invasions (2019-present)

Between 2014 and 2019, no new invasions on any islands were reported. In 2019, Arasamuthu et al., (2019) noted new establishment on two islands (Valai and Thalaiyari) in the third island group, without presence on the intervening islands, though these islands are also fringed with reefs (Panel 2, Figure 1). Many authors (Edward and J.R Bhatt 2012; Kamalakannan et al. 2014; Arasamuthu A. et al. 2023) suggested that *K. alvarezii* preferentially colonizes the branching coral *Acropora*, and proposed that an absence of *Acropora* could influence geographic spread.

During this period of time, contradictory narratives on invasion extent emerged in the literature. Arasamuthu et al., (2023) does not report the disappearance of *K. alvarezii* from invaded sites in the first island group. Krishnan et al., (2021) reports disappearance from one reported site, and calls into question the veracity of the reported invasion on one island. At the time of our field work, the TNFD had confirmed the absence of *K. alvarezii* from all islands within the first island group.

### 3.2 Insights from interviews

Responses for when collectors had last seen *K. alvarezii* ranged from 2-10 years prior to the interviews. There were three reported locations from the interview data.

The first (K3, Panel 2, Figure 1) is a channel between the mainland and Kurusudai Island, which experiences a great degree of boating pressure. It is also fairly close to the Pamban pass, where propagules of *K. alvarezii* are presumed to enter the Gulf of Mannar region. K3 is also the most accessible reef, and likely the most visited by seaweed collectors. Participants reported presence in K3 from ten years ago to two years ago.

The next two sites (K2 and K1, Panel 2, Figure 1) were difficult to separate from interview data due to their spatial proximity, but aligned with what has been reported in published literature. In K2 and K1, responses for when participants had last seen *K. alvarezii* ranged from 6 to 3 years prior.

Participants also referenced key events to triangulate remembered time periods. These include the Global Bleaching Event of 2016, and algal blooms in 2019, 2020 and 2021 (Raj et al. 2021). Consolidating information from interviews and published reports, we propose the following timeline of invasion history (Panel 1, Figure 1).

K1 and K2 had the longest durations of invasion, with presence reported annually or biannually from 2005-2016. One respondent suggested that *K. alvarezii* was observed in 2022, but this is not corroborated by the TNFD reports and gray literature. K3 likely had a duration of invasion of approximately eight years (2014-2022), and was also the most recently invaded site in the first island group. There was likely *K. alvarezii* presence as recently as 2022, but not after this. S2 was only described as invaded in a single short report in 2010 (J. K. Patterson Edward and J.R Bhatt 2012). No other independent survey or any interview data corroborates this. S1 likely had the shortest duration of invasion, of approximately 6 years, from 2014-2020. Its exact date of disappearance could not be estimated from interview data.

### 3.3 Insights from ecological surveys

Our surveys confirm a sharp decline in the extent of invasion. *K. alvarezii* presence was not recorded on any reef in the first island group, including the specific sites K1, K2, K3, S1 and S2, (Panel 2, Figure 1) and all proximate reefs.

We subsequently used a photo-quadrat method to evaluate cover at invaded sites (V1 and T1, Panel 2, Figure 1) and confirm absence in formerly invaded sites (K1, K2, K3, S1 and S2). In invaded sites, *K. alvarezii* cover was less than 1%, as derived from our benthic photo quadrats. *K. alvarezii* was observed in only two out of nine sites, and with very low cover.

Our ecological surveys, interviews and the literature review revealed that there are no new invaded sites, with the last new site being reported in 2019 (Arasamuthu A. et al. 2023). There is also a decline in *K. alvarezii* cover at the two invaded sites (T1 and V1). There is also no evidence, at present, of spread from invaded sites.

## 4. Discussion

Our interviews and ecological surveys, when considered in conjunction, provide strong evidence for the decline of *K. alvarezii* in the Gulf of Mannar, India. From the interviews, it appears that the first decline in presence at an invaded site occurred about a decade prior (2014-2016). Another period of decline took place between 6-9 years ago (2017-2019). It is likely that the first island group no longer had any *K. alvarezii* after 2022, which was confirmed by our ecological surveys. We also found no new invaded sites. This represents a significant decline in the extent of invasion, and is a novel finding in understanding the trajectory of *K. alvarezii* invasion.

### Causes of Decline

The TNFD has conducted annual manual removal drives since 2009, where free divers who regularly collect seaweed are employed to remove *K. alvarezii* from the reef by hand. Although manual removal efforts continue and do appear to have an effect on *K. alvarezii* abundance, it is unlikely to be the primary driving force behind its decline (Kamalakannan et al. 2014). Previous studies examined *K. alvarezii* cover post manual removal, and found that even with removal the cover remained at 0.3-8% (Kamalakannan et al. 2014). In fact, they suggested that manual removal may be actively harmful and promote the spread of propagules. This estimate of cover is far greater than our estimate of 0% in one invaded site (V1) and 0.04% in the second invaded site (T1) (Figure 2). Furthermore, many areas of the reef remain inaccessible for manual removal by free diving. *K. alvarezii* also often persists within interlocking branches of *Acropora*, which are also inaccessible for removal, underscoring the limitations of this management intervention. *K. alvarezii* cover was also found to attain or even exceed initial levels 6 months post removal (Kamalakannan et al. 2014), and our surveys were conducted prior to the annual removal drives. Larger-scale manual removal programmes were also not found to be successful in Hawaii (Winston et al. 2023). We did not document any corals being covered completely by *K. alvarezii*, nor did we observe any *K. alvarezii* induced mortality.

**Fig. 2.**
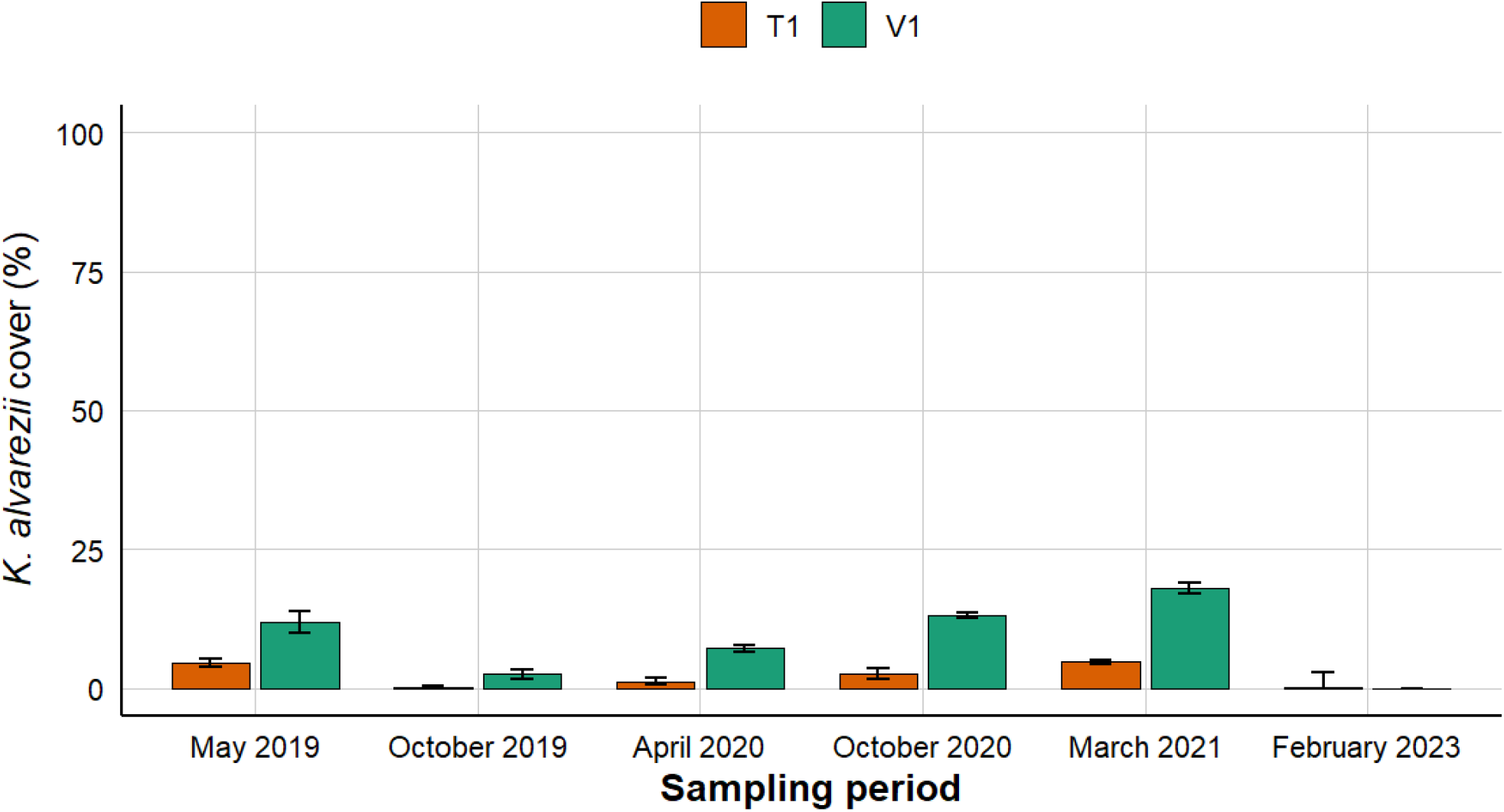
Change in cover of *K. alvarezii*. Data for May 2019, October 2019, April 2020, October 2020 and March 2021 were derived from Arasamuthu et al., (2023), and data from February 2023 was collected from our ecological surveys.

The proposed timeline of disappearance (Panel 2, Figure 1) suggests a possible link to climatic events in the region. 2014-2016 was a period of extreme heating in the region, with Sea Surface Temperatures (SSTs) exceeding 29ºC in April 2015 (Krishnan et al. 2018; Raj et al. 2021). Prolonged periods of high SSTs place a high degree of stress on shallow reef ecosystems, as is characteristic in the Gulf of Mannar (Raj et al. 2021). Raj et al. (2021) also note the greatest decrease in live coral cover in the first island group. This high temperature and loss in live coral cover could have, directly or indirectly, contributed to *K. alvarezii* decline by three mechanisms.

1. **A loss in live coral cover**. Coral cover in the Gulf of Mannar has declined considerably due to bleaching events (Krishnan et al. 2018; Raj et al. 2021). This may have led to a reduction in live coral substrate available for *K. alvarezii* to colonize, leading to a decline in *K. alvarezii* cover. *K. alvarezii* is also linked most frequently to live coral substrates, and has not been documented on dead corals (Kamalakannan et al. 2014; ArasamuthuA et al. 2023). We noted the same during our surveys.
2. **Direct heating effects on *K. alvarezii***. *K. alvarezii* is not a thermally tolerant species (Mantri et al. 2017). If temperatures exceed 32 ºC, as was observed in the Gulf of Mannar, *K. alvarezii* develops “ice-ice” disease. This is a whitening of the thalli that typically leads to mortality of the whole colony (Mantri et al. 2017). Direct heating effects may have caused large-scale mortality of *K. alvarezii*.
3. **Heating and heightened herbivory**. High SST events are known to increase herbivore metabolism (Winston et al. 2023), potentially leading them to consume novel algal resources. Where *K. alvarezii* does persist, it is between inaccessible, interlocking branches of *Acropora* colonies. These may act as a natural defense against herbivory, allowing some persistence of *K. alvarezii*. It also lends some evidence to the idea that *K. alvarezii* abundance is linked to *Acropora* cover (Edward and Bhatt 2012; Kamalakannan et al. 2014; Arasamuthu et al. 2023). Rather than *Acropora* being an inherently more suitable substrate for *K. alvarezii* growth, it is possible that *K. alvarezii* persists within branching corals as it is inaccessible to herbivores.

Another possibility is an overall loss in vitality of the *K. alvarezii* cultivar. Anecdotal evidence from farmers suggests that yield and vitality has declined substantially (YS personal communication, April 2024). It is possible that the trajectory in the wild mirrors that of cultivation. *K. alvarezii* reproduces asexually, and new genetic material has not been brought to India since the species’ introduction in 2002 (Mantri et al. 2017). All present cultivars descend from this initial population. This absence of genetic diversity might be leading to a loss in yield and vitality (Mantri et al. 2017, current conservation article).

It is also interesting to note that the decline of invasion in the Gulf of Mannar is paralleled by a decline in the other invaded regions of Hawaii and Panama. There was also a singular report from Costa Rica (Cabrera et al. 2019), but subsequent reports did not emerge from the region. In Panama, the invasion lasted only three years, from 2013-2016, and the decline was linked to urchin herbivory (Albright et al. 2022). In Hawaii, the invasion spanned almost 40 years, (Rodgers and Cox 1999; Woo 2000; Smith et al. 2002; Conklin and Smith 2005), with a sharp decline in 2013-2014. This decline was linked to flash floods on the island, which lowered the salinity of the bay, creating conditions that *K. alvarezii* could not tolerate (Winston et al. 2023). Exploring whether there is a common cause of decline, climatic or otherwise, would be an important line of inquiry.

It is possible that current observations and declines may be a smaller step in a larger boom and bust cycle, or a temporary decline, as invasive species have been known to follow these patterns (Strayer et al. 2017; Winston et al. 2023). The information presented in this report may represent a “bust”. Busts can occur due to self-limitation through resource depletion, the accumulation of specialised native enemies (like herbivores) or changes in environmental conditions (like warming).

### Implications of *K. alvarezii* decline

In the first island group, sites we assessed were dominated by dead coral. Unless the coming years are free from bleaching events, the ability of coral juveniles to use the existing substrate successfully and build new reefs remains limited. At invaded sites, a continued decline of *K. alvarezii* cover *could* improve reef health. Nonetheless, we noted that these invaded sites had the highest live coral cover and fish biomass. It is unlikely that this link is causal, nor does *K. alvarezii* offer any bleaching resistance, as *K. alvarezii* has been reported to cause significant damage to reefs in other regions (Woo 2000; Sellers et al. 2014). Rather, sites which continue to have *K. alvarezii* happened to maintain their live coral cover by escaping the most significant effects of the 2016 bleaching event.

The decline in invasion of *K. alvarezii* is paralleled by the decline in yield for farmers. Although the full extent of impacts is not explored here, the susceptibility of *K. alvarezii* to warming represents a vulnerability of the industry to climate change, and loss of livelihoods for farmers. Although aquaculture crops like *K. alvarezii* have brought significant livelihood gains and economic benefits in the past, it is vital to keep evaluating livelihood interventions as environmental conditions change.

### Short-Term Gains, Long-Term Consequences

Aquaculture is a significant pathway for novel biological invasions (Oficialdegui et al. 2025), which take place primarily through the translocation of cultivated species. The same traits that make a species profitable in an aquaculture context can make it more likely to invade. Aquaculture species are selected particularly for stress-tolerance, rapid reproduction and growth. Aquaculture can also contribute to the accidental release of species from farming operations, or the movement of associated organisms attached to cultured stock, equipment, or vessels (Namboothri et al. 2012). Marine aquaculture may indeed support food security and climate adaptation objectives, but the risk of introducing potentially invasive species will rise. This is a particular concern where there are no strong legislative protocols or quarantine measures for mitigating risk.

Climate change, the increasing frequency of El Niño warming events, and cyclones underscore the need to treat non-native aquaculture species with even greater caution. Literature indicates that climate change and warming can accelerate invasions, as disturbed ecosystems may be more vulnerable to non-native species (Mainka and Howard 2010). In marine systems, climate change and ocean warming are known to drive the process of tropicalisation, where tropically-adapted species are able to colonise higher and lower latitudes (Angeles-Gonzalez et al. 2026). More research is needed to understand how these two stressors interact, as interactions can be unpredictable and have unexpected consequences.

In addition to the ecological impacts of cultivating non-native species, it is also of merit to critically examine purported benefits of economic upliftment, carbon sequestration, and the “minimal” environmental impacts of seaweed farming, *K. alvarezii* yields have declined significantly under climate change (Largo et al. 2017; Kumar et al. 2020). Although aquaculture crops like *K. alvarezii* initially brought significant livelihood gains and economic benefits, these dynamics have changed in the context of ocean warming. Blue Economy interventions aim for the sustainable use of ocean resources, but sustainability has multiple dimensions. It is vital to carefully consider Blue Economy schemes, particularly those that hinge on non-native species introductions.

## Limitations

We were unable to infer long-term ecological impacts of this invasive alga due to the absence of long-term, publicly available data for the Gulf of Mannar. This also limited our ability to link the decline with a particular cause.

Consequently, we are missing information on the drivers of decline for one of the three regions globally where this commercially significant species has become invasive with a subsequent decline. The current study used interviews to bridge this gap, providing us with vital information on the time periods of decline. Our interview-based approach required participants to recall events from the past, likely resulting in somewhat imprecise timelines.

## Conclusion

Long term monitoring data is vital to situate invasive decline within the broader context of invasion ecology research. We recommend continuous monitoring, primarily through the collection of primary ecological data, supplemented by interviews with seaweed collectors. We also recommend a control-treatment-based approach to manual removal interventions to ascertain the true impact of removal, and to understand whether this management strategy needs to be modified in the context of the current decline.

It is vital to carefully consider the socio-ecological landscapes to which Blue Economy schemes are being introduced. The Gulf of Mannar, like many tropical and sub-tropical regions around the world, is extremely vulnerable to the impacts of climate change. Although non-native species and aquaculture may offer short-term gains, blue economy schemes can have long-term socioecological impacts and hidden costs for increasingly vulnerable ecosystems.

## 6. Statements and Declarations

### 6.3 Author Contributions

ED, SP, NN and YS contributed to the study conception and design. ED, SP, NN and YS contributed to material preparation, data collection and analysis. The first draft of the manuscript was written by YS and all authors commented on subsequent versions of the manuscript. All authors read and approved the final manuscript and consent for publication.

### 6.4 Ethics approval and consent to participate

Animal and human ethics clearance were obtained from Nature Conservation Foundation, Mysuru [Number : NCF-EC-05/01/2024-(89)]. Principles of informed prior consent were applied during the interview process.

### 6.5 Data availability

Data can be made available on request to the author.

## 6.6 Acknowledgments

We thank Suresh Kumar, Praveen, Sarah Christin and Keerthikrutha Seetharaman who aided extensively in data collection and field surveys. Johnson contributed to experimental work. Joseph Anand provided a great deal of support. The Dakshin Foundation Field Station at Palk Bay provided significant logistic aid.

