## Supplementary for "Decline in *Kappaphycus alvarezii* invasion in the Gulf of Mannar, India"

**Supplementary 1: Complete Search Terms**

Search string 1: Invasion patterns globally:

("Kappaphycus alvarezii" OR Kappaphycus OR "Eucheuma cottonii" OR "Eucheuma denticulatum")
AND
(invasive OR invasion OR introduced OR non-native OR exotic OR spread OR colonization OR establishment)

Search string 2: Mechanisms of invasion:

("Kappaphycus alvarezii")
AND
(growth OR fragmentation OR dispersal OR "vegetative propagation" OR tolerance OR "phenotypic plasticity")
AND
(invasive OR spread OR establishment)

Search string 3: Management practices

("Kappaphycus" OR "Eucheuma")
AND
(removal OR control OR management OR mitigation OR biocontrol OR herbivory OR restoration)
AND
(coral OR reef)

Search string 4: Capturing all case studies globally

("Kappaphycus alvarezii")
AND
(invasive OR invasion OR introduced)
AND
(Hawaii OR India OR Panama OR Brazil OR Caribbean OR Pacific OR "Gulf of Mannar")

**Supplementary 2: Complete List of References**

13. Ask EI, Batibasaga A, Zertuche J, San M (2003) Three decades of Kappaphycus alvarezii (Rhodophyta) introduction to non-endemic locations. Proc Int Seaweed Symp 17:49–57
14. Ateweberhan M, Rougier A, Rakotomahazo C (2015) Influence of environmental factors and farming technique on growth and health of farmed Kappaphycus alvarezii (cottonii) in south-west Madagascar. J Appl Phycol 27:923–934. https://doi.org/10.1007/s10811-014-0378-3
15. Balasubramanian K, J JJJ, Nagendran A, et al (2014) Impact of removal of invasive species Kappaphycus alvarezii from coral reef ecosystem in Gulf of Mannar, India. Current science 106:25–2014
16. Bast F, John AA, Bhushan S (2016) Molecular assessment of invasive Carrageenophyte Kappaphycus alvarezii from India based on ITS-1 sequences. Webbia 71:287–292. https://doi.org/10.1080/00837792.2016.1221187
17. Becker A, Whitfield AK, Cowley PD, et al (2016) Tidal amplitude and fish abundance in the mouth region of a small estuary. J Fish Biol 89:1851–1856. https://doi.org/10.1111/jfb.13056
18. Bellwood DR Sleeping Functional Group Drives Coral-Reef Recovery. Current Biology
19. Bennett NJ, Cisneros-Montemayor AM, Blythe J, et al (2019) Towards a sustainable and equitable blue economy. Nat Sustain 2:991–993. https://doi.org/10.1038/s41893-019-0404-1
20. Bennett S, Vergés A, Bellwood DR (2010) Branching coral as a macroalgal refuge in a marginal coral reef system. Coral Reefs 29:471–480. https://doi.org/10.1007/s00338-010-0594-5
21. Bhatt J, JK Patterson Edward (2012) Impacts of cultivation of Kappaphycus alvarezii on coral reef environs of the Gulf of Mannar and Palk Bay, South-eastern India. pp 89–98
22. Bhuyan MdS (2023) Ecological risks associated with seaweed cultivation and identifying risk minimization approaches. Algal Research 69:102967. https://doi.org/10.1016/j.algal.2022.102967
23. Bonacchi A, Gasperini S, Bartolommei P, et al (2014) Seasonal Food Selection In Small Small Mammals: A Cafeteria Experiment
24. Brandl SJ, Rasher DB, Côté IM, et al (2019) Coral reef ecosystem functioning: eight core processes and the role of biodiversity. Frontiers in Ecology and the Environment 17:445–454. https://doi.org/10.1002/fee.2088
25. Brooks M E, Kristensen K, Benthem K J ,van, et al (2017) glmmTMB Balances Speed and Flexibility Among Packages for Zero-inflated Generalized Linear Mixed Modeling. The R Journal 9:378. https://doi.org/10.32614/RJ-2017-066
26. Budzałek G, Śliwińska-Wilczewska S, Wiśniewska K, et al (2021) Macroalgal Defense against Competitors and Herbivores. International Journal of Molecular Sciences 22:7865. https://doi.org/10.3390/ijms22157865
27. Bureau TH (2023) Union Minister lays foundation stone for seaweed park near Ramanathapuram. The Hindu

28. Buschmann AH, Camus C, Infante J, et al (2017) Seaweed production: overview of the global state of exploitation, farming and emerging research activity. European Journal of Phycology 52:391–406. https://doi.org/10.1080/09670262.2017.1365175
29. Cabrera R, Umanzor S, Díaz-Larrea J, Araújo PG (2019) Kappaphycus alvarezii (Rhodophyta): New Record of an Exotic Species for the Caribbean Coast of Costa Rica. American Journal of Plant Sciences 10:1888–1902. https://doi.org/10.4236/ajps.2019.1010133
30. Cai J, Lovatelli A, Aguilar-Manjarrez, J., et al (2021) Seaweeds and microalgae: an overview for unlocking their potential in global aquaculture development. Food and Agriculture Organization of the United Nations, Rome
31. Campbell R, Hotchkiss S (2017) Carrageenan Industry Market Overview. In: Hurtado AQ, Critchley AT, Neish IC (eds) Tropical Seaweed Farming Trends, Problems and Opportunities: Focus on Kappaphycus and Eucheuma of Commerce. Springer International Publishing, Cham, pp 193–205
32. Castelar B, de Siqueira MF, Sánchez-Tapia A, Reis RP (2015) Risk analysis using species distribution modeling to support public policies for the alien alga Kappaphycus alvarezii aquaculture in Brazil. Aquaculture 446:217–226. https://doi.org/10.1016/j.aquaculture.2015.05.012
33. Castro-Sanguino C, Lovelock C, Mumby PJ (2016) The effect of structurally complex corals and herbivory on the dynamics of Halimeda. Coral Reefs 35:597–609. https://doi.org/10.1007/s00338-016-1412-5
34. Chandrasekaran S, Nagendran A, Pandiaraja D, et al (2008) Bioinvasion of Kappaphycus alvarezii on corals in the Gulf of Mannar, India. Current Science 94:
35. Chapman D, Ranelletti M, Kaushik S (2006) Invasive Marine Algae: An Ecological Perspective. The Botanical Review 72:153–178. https://doi.org/10.1663/0006-8101(2006)72%255B153:IMAAEP%255D2.0.CO;2
36. Chong-Seng KM, Nash KL, Bellwood DR, Graham NAJ (2014) Macroalgal herbivory on recovering versus degrading coral reefs. Coral Reefs 33:409–419. https://doi.org/10.1007/s00338-014-1134-5
37. Cisneros-Montemayor AM, Moreno-Báez M, Reygondeau G, et al (2021) Enabling conditions for an equitable and sustainable blue economy. Nature 591:396–401. https://doi.org/10.1038/s41586-021-03327-3
38. Conklin EJ, Smith JE (2005) Abundance and Spread of the Invasive Red Algae, Kappaphycus spp., in Kane'ohe Bay, Hawai'i and an Experimental Assessment of Management Options. Biol Invasions 7:1029–1039. https://doi.org/10.1007/s10530-004-3125-x
39. Connell SD, Russell BD, Irving AD (2011) Can strong consumer and producer effects be reconciled to better forecast 'catastrophic' phase-shifts in marine ecosystems? Journal of

Experimental Marine Biology and Ecology 400:296–301. https://doi.org/10.1016/j.jembe.2011.02.031

40. Cooke RG (2004) Rich, poor, shaman, child: animals, rank, and status in the “Gran Code” culture area of pre-Columbian Panama. In: International Council for Archaeozoology, Jones S, Neer W van, Ervynck A (eds) Behaviour behind bones: the zooarchaeology of ritual, religion, status, and identity. Oxbow ; David Brown Book Co, Oxford : Oakville, CT
41. Côté IM, Smith NS (2018) The lionfish *Pterois* sp. invasion: Has the worst-case scenario come to pass? Journal of Fish Biology 92:660–689. https://doi.org/10.1111/jfb.13544
42. Couperus B, Gastauer S, Fässler SMM, et al (2016) Abundance and tidal behaviour of pelagic fish in the gateway to the Wadden Sea. Journal of Sea Research 109:42–51. https://doi.org/10.1016/j.seares.2016.01.007
43. Dahl AL (1973) Surface area in ecological analysis: Quantification of benthic coral-reef algae. Marine Biology 23:239–249. https://doi.org/10.1007/BF00389331
44. Darling ES, McClanahan TR, Maina J, et al (2019) Social–environmental drivers inform strategic management of coral reefs in the Anthropocene. Nat Ecol Evol 3:1341–1350. https://doi.org/10.1038/s41559-019-0953-8
45. Deb A (2019) Creating Opportunities for Women through Seaweed Farming. In: Harnessing Nature. https://harnessingnature.wordpress.com/2019/02/24/creating-opportunities-for-women-through-seaweed-farming/. Accessed 23 June 2023
46. Department of Fisheries (2022) Transforming Gisheries: PRadhan Mantri Matsya Sampadha Yojana. Department of Fisheries
47. Dillehay TD, Ramírez C, Pino M, et al (2008) Monte Verde: Seaweed, Food, Medicine, and the Peopling of South America. Science 320:784–786. https://doi.org/10.1126/science.1156533
48. Dolan AH, Walker IJ (2006) Understanding Vulnerability of Coastal Communities to Climate Change Related Risks. Journal of Coastal Research 1316–1323
49. Donovan MK, Friedlander AM, Lecky J, et al (2018) Combining fish and benthic communities into multiple regimes reveals complex reef dynamics. Sci Rep 8:16943. https://doi.org/10.1038/s41598-018-35057-4
50. Doty MS, Caddy JF, Doty MS, et al (eds) (1987) The Production and Use of Euchema. In: Case studies of seven commercial seaweed resources. Food and Agriculture Organization of the United Nations, Rome, pp 123–164
51. Du Y-Q, Jueterbock A, Firdaus M, et al (2023) Niche comparison and range shifts for two Kappaphycus species in the Indo-Pacific Ocean under climate change. Ecological Indicators 154:110900. https://doi.org/10.1016/j.ecolind.2023.110900
52. Duffy JE, Hay ME (1990) Seaweed Adaptations to Herbivory. BioScience 40:368–375. https://doi.org/10.2307/1311214
53. Edward JKP, Mathews G, Raj KD, et al (2012) Coral reefs of Gulf of Mannar, India - signs of resilience

92. Kamalakannan B, Jeevamani JJJ, Nagendran NA, et al (2010) Turbinaria sp. as victims to Kappaphycusalvarezii in reefs of Gulf of Mannar, India. Coral Reefs 29:1077–1077
93. Kamalakannan B, Jeevamani JJJ, Nagendran NA, et al (2014) Impact of removal of invasive species Kappaphycus alvarezii from coral reef ecosystem in Gulf of Mannar, India. Current Science 106:1401–1408
94. Kambey CSB, Sondak CFA, Chung I-K (2020) Potential growth and nutrient removal of Kappaphycus alvarezii in a fish floating-net cage system in Sekotong Bay, Lombok, Indonesia. Journal of the World Aquaculture Society 51:944–959. https://doi.org/10.1111/jwas.12683
95. Karthikeyan S, Kulandaisamy P, Saravanan P, et al (2022) Agriculture Drought Management in Ramanathapuram District of Tamil Nadu, India. 8:59–65. https://doi.org/10.3233/JCC220005
96. Kelkar N, Arthur R, Marbà N, Alcoverro T (2013) Greener pastures? High-density feeding aggregations of green turtles precipitate species shifts in seagrass meadows. Journal of Ecology 101:1158–1168. https://doi.org/10.1111/1365-2745.12122
97. Kelly E, Eynaud Y, Williams I, et al (2017) A budget of algal production and consumption by herbivorous fish in an herbivore fisheries management area, Maui, Hawaii. Ecosphere 8:e01899. https://doi.org/10.1002/ecs2.1899
98. Kelly ELA, Cannon AL, Smith JE (2020) Environmental impacts and implications of tropical carrageenophyte seaweed farming. Conservation Biology 34:326–337. https://doi.org/10.1111/cobi.13462
99. Kelly ELA, Eynaud Y, Clements SM, et al (2016) Investigating functional redundancy versus complementarity in Hawaiian herbivorous coral reef fishes. Oecologia 182:1151–1163. https://doi.org/10.1007/s00442-016-3724-0
100. Kerr JNQ, Paul VJ (1995) Animal-plant defense association: the soft coral Sinularia sp. (Cnidaria, Alcyonacea) protects Halimeda spp. from herbivory. Journal of Experimental Marine Biology and Ecology 186:183–205. https://doi.org/10.1016/0022-0981(94)00153-5
101. Klein J, Verlaque M (2008) The *Caulerpa racemosa* invasion: A critical review. Marine Pollution Bulletin 56:205–225. https://doi.org/10.1016/j.marpolbul.2007.09.043
102. Krimou S, Gairin E, Gautrand L, et al (2023) Herbivory effects of sea urchin species on a coral reef (Bora-Bora, French Polynesia). Journal of Experimental Marine Biology and Ecology 564:151900. https://doi.org/10.1016/j.jembe.2023.151900
103. Krishnamurthy R Coral breach: A silent, catastrophic invasion has happened in the Gulf of Mannar; here is how. Down to Earth
104. Krishnan P, Abhilash KR, Sreeraj CR, et al (2021) Balancing livelihood enhancement and ecosystem conservation in seaweed farmed areas: A case study from Gulf of Mannar Biosphere Reserve, India. Ocean & Coastal Management 207:105590. https://doi.org/10.1016/j.ocecoaman.2021.105590

118. Mack RN, Simberloff D, Mark Lonsdale W, et al (2000) Biotic Invasions: Causes, Epidemiology, Global Consequences, and Control. Ecological Applications 10:689–710. https://doi.org/10.1890/1051-0761(2000)010%255B0689:BICEGC%255D2.0.CO;2
119. Mainka SA, Howard GW (2010) Climate change and invasive species: double jeopardy. Integrative Zoology 5:102–111. https://doi.org/10.1111/j.1749-4877.2010.00193.x
120. Mandal GS, Mohapatra M (2010) Cyclone Hazard Prone Districts of India : A Report
121. Mandal S, Mantri V, Haldar S, et al (2010) Invasion potential of Kappaphycus alvarezii on corals at Kurusadai Island, Gulf of Mannar, India. International Journal on Algae 25:205–216
122. Mantri V, Eswaran K, Munisamy S, et al (2017a) An appraisal on commercial farming of Kappaphycus alvareziiin India: success in diversification of livelihood and prospects. Journal of Applied Phycology 29:. https://doi.org/10.1007/s10811-016-0948-7
123. Mantri VA, Ashok KS, Musamil TM, et al (2017b) Tube-net farming and device for efficient tissue segregation for industrially important agarophyte Gracilaria edulis (Rhodophyta). Aquacultural Engineering 77:132–135. https://doi.org/10.1016/j.aquaeng.2017.04.003
124. Mantri VA, Kavale MG, Kazi MA (2020) Seaweed Biodiversity of India: Reviewing Current Knowledge to Identify Gaps, Challenges, and Opportunities. Diversity 12:13. https://doi.org/10.3390/d12010013
125. Mantyka CS, Bellwood DR (2007) Direct evaluation of macroalgal removal by herbivorous coral reef fishes. Coral Reefs 26:435–442. https://doi.org/10.1007/s00338-007-0214-1
126. McCook L, Jompa J, Diaz-Pulido G (2001) Competition between corals and algae on coral reefs: a review of evidence and mechanisms. Coral Reefs 19:400–417. https://doi.org/10.1007/s003380000129
127. McCook LJ (1999) Macroalgae, nutrients and phase shifts on coral reefs: scientific issues and management consequences for the Great Barrier Reef. Coral Reefs 18:357–367. https://doi.org/10.1007/s003380050213
128. McManus JW, Polsenberg JF (2004) Coral–algal phase shifts on coral reefs: Ecological and environmental aspects. Progress in Oceanography 60:263–279. https://doi.org/10.1016/j.pocean.2004.02.014
129. McPhaden MJ (2015) Playing hide and seek with El Niño. Nature Clim Change 5:791–795. https://doi.org/10.1038/nclimate2775
130. Meenakshisundaram G, Thiruppathi S, Sahu N, et al (2006) In situ observations on preferential grazing of seaweeds by some herbivores. Current Science 91:1256–1260
131. Mercado JM, Gómez-Jakobsen F, Korbee N, et al (2022) Analyzing environmental factors that favor the growth of the invasive brown macroalga *Rugulopteryx okamurae* (Ochrophyta): The probable role of the nutrient excess. Marine Pollution Bulletin 174:113315. https://doi.org/10.1016/j.marpolbul.2021.113315

132. Mondal M (2023) The Rise and Fall of a Commercial Seaweed – and a Community's Fortunes With It. The Wire Scince
133. Monteiro CA, Engelen AH, Santos ROP (2009) Macro- and mesoherbivores prefer native seaweeds over the invasive brown seaweed Sargassum muticum: a potential regulating role on invasions. Mar Biol 156:2505–2515. https://doi.org/10.1007/s00227-009-1275-1
134. Mouritsen OG, Rhatigan P, Cornish ML, et al (2021) Saved by seaweeds: phyconomic contributions in times of crises. J Appl Phycol 33:443–458. https://doi.org/10.1007/s10811-020-02256-4
135. Msuya F (2006) The Impact of Seaweed Farming on the Social and Economic Structure of Seaweed Farming Communities in Zanzibar, Tanzania. In: World seaweed resources: an authoritative reference system. ETI Bioinformatics, Amsterdam
136. Msuya F (2010) Development of seaweed cultivation in Tanzania: the role of the University of Dar es Salaam and other institutions. Aquaculture Compendium
137. Msuya F (2011) Environmental changes and their impact on seaweed farming in Tanzania. World Aquaculture World Aquaculture:34-37,71
138. Msuya FE (2020) Seaweed resources of Tanzania: status, potential species, challenges and development potentials. Botanica Marina 63:371–380. https://doi.org/10.1515/bot-2019-0056
139. Mulyani S, Tuwo A, Syamsuddin R, Jompa J (2018) Effect of seaweed Kappaphycus alvarezii aquaculture on growth and survival of coral Acropora muricata
140. Muralidharan R, Rai ND (2020) Violent maritime spaces: Conservation and security in Gulf of Mannar Marine National Park, India. Political Geography 80:102160. https://doi.org/10.1016/j.polgeo.2020.102160
141. Myers SS, Gaffikin L, Golden CD, et al (2013) Human health impacts of ecosystem alteration. Proceedings of the National Academy of Sciences 110:18753–18760. https://doi.org/10.1073/pnas.1218656110
142. Nagaraj A (2020) India's women seaweed divers swim against the tide of climate change. Reuters
143. Naveen Namboothri, Rauf Ali, Ankila Hiremath (2012) BIOLOGICAL INVASIONS OF MARINE ECOSYSTEMS: Concerns for Tropical Nations
144. Neill PE, Alcalde O, Faugeron S, et al (2006) Invasion of *Codium fragile* ssp. *tomentosoides* in northern Chile: A new threat for *Gracilaria* farming. Aquaculture 259:202–210. https://doi.org/10.1016/j.aquaculture.2006.05.009
145. Neilson BJ, Wall CB, Mancini FT, Gewecke CA (2018) Herbivore biocontrol and manual removal successfully reduce invasive macroalgae on coral reefs. PeerJ 6:e5332. https://doi.org/10.7717/peerj.5332

146. NOAA (2024) 2023 was the warmest year in the modern temperature record. http://www.climate.gov/news-features/featured-images/2023-was-warmest-year-modern-temperature-record. Accessed 20 July 2024
147. Oficialdegui FJ, Soto I, Balzani P, et al (2025) Non-Native Species in Aquaculture: Burgeoning Production and Environmental Sustainability Risks. Reviews in Aquaculture 17:e70037. https://doi.org/10.1111/raq.70037
148. O'Riordan T (2019) Avoiding an Unjust Transition. Environment: Science and Policy for Sustainable Development 61:2–3. https://doi.org/10.1080/00139157.2019.1564211
149. Pagès JF, Smith TM, Tomas F, et al (2018) Contrasting effects of ocean warming on different components of plant-herbivore interactions. Marine Pollution Bulletin 134:55–65. https://doi.org/10.1016/j.marpolbul.2017.10.036
150. Palani Kumar (2019) Tamil Nadu's seaweed harvesters in rough seas. In: People's Archive of Rural India. https://ruralindiaonline.org/en/articles/tamil-nadus-seaweed-harvesters-in-rough-seas/. Accessed 23 June 2023
151. Pandiaraja D, Nagendran NA, Murugeswari D, Mishra VN (2017) Spatial competition mathematical model analysis for the invasion, removal of Kappaphycus Algae in gulf of Mannar with propagation delays. Commun Math Biol Neurosci 2017:Article ID 10. https://doi.org/10.28919/cmbn/3317
152. Panippiḷḷa R, Mariarajan T (2014) A participatory study of the traditional knowledge of fishing communities in the Gulf of Mannar, India: the communities of Chinnapalam and Bharathi Nagar, Ramanathapuram district, Tamil Nadu, India. International Collective in Support of Fishworkers, Chennai, India
153. Pereira N, Velecar XN (2005) Is Gulf of Mannar heading for marine bioinvasion? Current Science 89:1309–1310
154. Pérez-Harguindeguy N, Díaz S, Vendramini F, et al (2003) Leaf traits and herbivore selection in the field and in cafeteria experiments. Austral Ecology 28:642–650. https://doi.org/10.1046/j.1442-9993.2003.01321.x
155. Periyasamy C, Anantharaman P, Balasubramanian T, Rao PVS (2014) Seasonal variation in growth and carrageenan yield in cultivated Kappaphycus alvarezii (Doty) Doty on the coastal waters of Ramanathapuram district, Tamil Nadu. J Appl Phycol 26:803–810. https://doi.org/10.1007/s10811-014-0256-z
156. Poray AK, Carpenter RC (2014) Distributions of coral reef macroalgae in a back reef habitat in Moorea, French Polynesia. Coral Reefs 33:67–76. https://doi.org/10.1007/s00338-013-1104-3
157. Prasad Behera D, Vadodariya V, Veeragurunathan V, et al (2022) Seaweeds cultivation methods and their role in climate mitigation and environmental cleanup. Total Environment Research Themes 3–4:100016. https://doi.org/10.1016/j.totert.2022.100016

158. Provan J, Murphy S, Maggs CA (2005) Tracking the invasive history of the green alga Codium fragile ssp. tomentosoides. Molecular Ecology 14:189–194. https://doi.org/10.1111/j.1365-294X.2004.02384.x
159. Puk LD, Marshell A, Dwyer J, et al (2020) Refuge-dependent herbivory controls a key macroalga on coral reefs. Coral Reefs 39:953–965. https://doi.org/10.1007/s00338-020-01915-9
160. Raj KD, Aeby GS, Mathews G, et al (2021) Coral reef resilience differs among islands within the Gulf of Mannar, southeast India, following successive coral bleaching events. Coral Reefs 40:1029–1044. https://doi.org/10.1007/s00338-021-02102-0
161. Raj KD, Mathews G, Obura DO, et al (2020) Low oxygen levels caused by Noctiluca scintillans bloom kills corals in Gulf of Mannar, India. Sci Rep 10:22133. https://doi.org/10.1038/s41598-020-79152-x
162. Rajagopalan R (2015) Q & A: Lakshmi Murthy. In: International Collective in Support of Fishworkers. https://www.icsf.net/yemaya/q-a-22/. Accessed 23 June 2023
163. Rameshkumar. P, Thirumalaiselvan PS, Raman M, et al (2023) Monitoring of Harmful Algal Bloom (HAB) of *Noctiluca scintillans* (Macartney) along the Gulf of Mannar, India using in-situ and satellite observations and its impact on wild and maricultured finfishes. Marine Pollution Bulletin 188:114611. https://doi.org/10.1016/j.marpolbul.2023.114611
164. Reimann L, Vafeidis AT, Honsel LE (2023) Population development as a driver of coastal risk: Current trends and future pathways. Cambridge Prisms: Coastal Futures 1:e14. https://doi.org/10.1017/cft.2023.3
165. Reis R (2000) Invasive potential of Kappaphycus alvarezii off the south coast of Rio de Janeiro state, Brazil: a contribution to environmentally secure cultivation in the tropics. Botanica Marina
166. Robert Panipilla, Marirajan T (2014) A Participatory Study of the Traditional Knowledge of Fishing Communities in the Gulf of Mannar, India
167. Rodgers S, Cox EF (1999) Rate of Spread of Introduced Rhodophytes Kappaphycus alvarezii, Kappaphycus striatum, and Gracilaria salicornia and Their Current Distribution in Kane'ohe Bay, O'ahu Hawai'i. Pacific Science 53:232–241
168. Roff G (2021) Evolutionary History Drives Biogeographic Patterns of Coral Reef Resilience. BioScience 71:26–39. https://doi.org/10.1093/biosci/biaa145
169. Russell DJ (1983) Ecology of the Imported Red Seaweed Eucheuma striatum Schmitz on Coconut Island, Oahu, Hawaii. Pacific Science 37:87–107
170. Santamaría J, Golo R, Verdura J, et al (2022) Learning takes time: Biotic resistance by native herbivores increases through the invasion process. Ecology Letters 25:2525–2539. https://doi.org/10.1111/ele.14115

171. Santoso A, Mcphaden MJ, Cai W (2017) The Defining Characteristics of ENSO Extremes and the Strong 2015/2016 El Niño. Reviews of Geophysics 55:1079–1129. https://doi.org/10.1002/2017RG000560
172. Saravanan R, Sadiq IS, Jawahar P (2018) Sea urchin diversity and its resources from the Gulf of Mannar. Centre for Innovation in Science and Social Action, Thiruvananthapuram, pp 194–196
173. Schaffelke B, Smith JE, Hewitt CL (2006) Introduced Macroalgae – a Growing Concern. J Appl Phycol 18:529–541. https://doi.org/10.1007/s10811-006-9074-2
174. Seebens H, Gastner MT, Blasius B (2013) The risk of marine bioinvasion caused by global shipping. Ecology Letters 16:782–790. https://doi.org/10.1111/ele.12111
175. Sellers A, Saltonstall K, Davidson T (2014) The introduced alga Kappaphycus alvarezii (Doty ex PC Silva, 1996) in abandoned cultivation sites in Bocas del Toro, Panama. BioInvasions Records 4:. http://dx.doi.org/10.3391/bir.2015.4.1.01
176. Senthalir S (2019) These Tamil Nadu women have been diving deep for seaweed – but now they are deeply worried. In: Scroll.in. https://scroll.in/article/913591/these-tamil-nadu-women-seaweed-divers-fear-the-loss-of-their-only-source-of-livelihood. Accessed 23 June 2023
177. Shigesada N, Kawasaki K (1997) Biological Invasions: Theory and Practice. Oxford University Press, UK
178. Sievanen L, Crawford B, Pollnac R, Lowe C (2005) Weeding through assumptions of livelihood approaches in ICM: Seaweed farming in the Philippines and Indonesia. Ocean & Coastal Management 48:297–313. https://doi.org/10.1016/j.ocecoaman.2005.04.015
179. Silver JJ, Gray NJ, Campbell LM, et al (2015) Blue Economy and Competing Discourses in International Oceans Governance. The Journal of Environment & Development 24:135–160. https://doi.org/10.1177/1070496515580797
180. Simberloff D, Gibbons L (2004) Now you See them, Now you don't! – Population Crashes of Established Introduced Species. Biological Invasions 6:161–172. https://doi.org/10.1023/B:BINV.0000022133.49752.46
181. Singh BS, Kaur I, Tuteja M, et al (2021) SEAWEED FARMING ENTREPRENEURSHIP: Recommendations Emerging From International Workshop On ENTREPRENEURSHIP DEVELOPMENT THROUGH SEAWEED BUSINESS BY COOPERATIVES. Laxmanrao Inamdar National Academy for Cooperative Research and Development (LINAC
182. Smith JE (2003) 153 Invasive Macroalgae on Tropical Reefs: Impacts, Interactions, Mechanisms and Management. Journal of Phycology 39:53–53. https://doi.org/10.1111/j.0022-3646.2003.03906001_153.x
183. Smith JE, Hunter CL, Smith CM (2002) Distribution and Reproductive Characteristics of Nonindigenous and Invasive Marine Algae in the Hawaiian Islands. Pacific Science 56:299–315

197. Thomsen MS, Wernberg T, South PM, Schiel DR (2016) Non-native Seaweeds Drive Changes in Marine Coastal Communities Around the World. In: Hu Z-M, Fraser C (eds)

Seaweed Phylogeography: Adaptation and Evolution of Seaweeds under Environmental Change. Springer Netherlands, Dordrecht, pp 147–185

207. Walter Scott (2014) Ramnad: paradoxes of a backward constituency. The Hindu

208. Winston M, Fuller K, Neilson BJ, Donovan MK (2023) Complex drivers of invasive macroalgae boom and bust in Kāne'ohe Bay, Hawai'i. Marine Pollution Bulletin 197:115744. https://doi.org/10.1016/j.marpolbul.2023.115744

209. Woo MML (2000) Ecological Impacts and Interactions of the Introduced Red Alga, Kappaphycus Striatum, In Kane'OHE Bay, O'Ahu. University of Hawaii

210. Wright JH, Hill NAO, Roe D, et al (2016) Reframing the concept of alternative livelihoods. Conserv Biol 30:7–13. https://doi.org/10.1111/cobi.12607
211. Zuccarello GC, Critchley AT, Smith J, et al (2006) Systematics and genetic variation in commercial shape Kappaphycus and shape Eucheuma (Solieriaceae, Rhodophyta). J Appl Phycol 18:643–651. https://doi.org/10.1007/s10811-006-9066-2
212. Zuniga-Jara S, Marin-Riffo M (2016) Bioeconomic analysis of small-scale cultures of Kappaphycus alvarezii (Doty) Doty in India. J Appl Phycol 28:1133–1143. https://doi.org/10.1007/s10811-015-0616-3
213. Global seafood market value forecast 2028. In: Statista. https://www.statista.com/statistics/821023/global-seafood-market-value/. Accessed 19 July 2024a
214. Gulf of Mannar Marine Biosphere Reserve | Ramsar Sites Information Service. https://rsis.ramsar.org/ris/2472. Accessed 23 July 2024b
215. (2020) Cyclonic Storm, "BUREVI‟ over the Bay of Bengal (30th November - 05th December 2020): A Report

**Supplementary 3: Interview Questions and Consent Form in English and Tamil**

### Interview Questions (English)

1. In what places have you encountered Kappaphycus?
2. When did you last see it in those areas?
3. Do you recall any large changes or climate related events occurring in the last few years? (e.g, algal blooms, coral bleaching, unusually warm water) If so, what happened, and when?
4. When you observed Kappaphycus, what corals was it growing on?
5. Have you ever heard of or seen any fish feeding on Kappaphycus?
6. Have you seen any regular patterns in growth of Kappaphycus? For example, is there a particular season in which abundances are very high?
7. Has Kappaphycus had any negative impacts on your seaweed collection work? For example, decreased amount of algae you can harvest?
8. In your opinion, is this algae good for sea-life or bad for sea-life?
9. In your opinion, was it right to ban cultivation in the Gulf of Mannar? Has the ban negatively affected livelihoods in your region?

### Interview Questions (Tamil)

1.பெப்சி பாசியை எங்கே பார்த்தீரகள்?

2.எந்த இடத்தில் எப்பொழுது பார்த்தீர்கள்?

3.எந்தெந்த வருடங்களில் பருவநிலை மாற்றங்கள் ஏற்பட்டது?அப்படி கடலில் ஏற்ப்பட்ட மாற்றம் என்ன?(பச்சைநீர் ஓட்டம் போன்றவை)

4.இந்த பாசி எந்த பாறைகளில் மீதெல்லாம் வளர்கிறது?

5.இந்த பாசியை உண்ணக்கூடிய மீன்கள் என்னென்ன? அவைகளை பர்த்துள்ளீர்களா?

6.இந்த பெப்சி பாசியினால் நீங்கள் செய்யும் தொழிலுக்கு ஏதேனும் பிரச்சனை வந்ததா? (எ.க: நீங்கள் சேகரிக்கும் பாசியின் அளவு அதன் வளர்ச்சி நிலைகளில் ஏதேனும் பாதிப்பு ஏற்பட்டடுள்ளதா?)

என்ன விதமான பாதிப்பு எற்ப்பட்டுள்ளது?

7.எவ்விதமான மீன்கள் இந்த பாசியை உண்கின்றன?

8.உங்களுடைய கருதுப்படி இந்த பாசி கடலுக்கு நன்மை செய்கிறதா அல்லது தீமை செய்கிறதா?

9.தெற்க்கு பகுதியில் இத்தொழிலினை தடை செய்வது சரியா அல்லது உங்களின் தொழில் முன்னேற்றத்திற்குத் தேவையா?

### Interview Consent Form (English)

**Research Project Title:** *Examining the ecological interactions and impacts of the cultivated algae Kappaphycus alvarezii*

**Investigators:**
Yamini Srikanth, National Centre for Biological Sciences (, 9886901322)
Dr. Elrika D'Souza, Nature Conservation Foundation
Dr. Sandeep Pulla,
Dr. Naveen Namboothri, Dakshin Foundation

**Nature and Purpose of the Project:**

*Kappaphycus alvarezii* is an economically important algae cultivated in the Palk Bay. Concerns have arisen about its negative effects on coral reefs. We aim to understand similarities and differences in sites where it has been found in the Gulf of Mannar. We also want to understand herbivory.

**Nature and Purpose of the Interview:**

*Kappaphycus* has disappeared from all previously invaded sites across Kurusudai and Shingle islands. Seaweed collectors often visit these regions more frequently from scientists, and have an in-depth understanding of how these reefs have changed over the last few years. We want to understand when *Kappaphycus* disappeared from Kurusudai and Shingle, and evaluate potential causes.

The interview will take thirty to forty-five minutes. We don't anticipate that there are any risks associated with your participation, but you have the right to stop the interview or withdraw from the research at any time.

Thank you for agreeing to be interviewed as part of the above research project. Ethical procedures for academic research require that interviewees explicitly agree to being interviewed and how the information contained in their interview will be used. This consent form is necessary for us to ensure that you understand the purpose of your involvement and that you agree to the conditions of your participation.

**Consent to Take Part in Research:**

- I, ________________________________ voluntarily agree to participate in this research study
- I understand that even if I agree to participate now, I can withdraw at any time or refuse to answer any question without any consequences of any kind
- I understand that I can withdraw permission to use data from my interview by contacting Yamini Srikanth within two weeks after the interview, in which case the material will be deleted.
- I have had the purpose and nature of the study explained to me in writing or verbally and I have had the opportunity to ask questions about the study
- I understand that I will not benefit directly from participating in this research.
- I agree to my interview being audio-recorded.
- I understand that all information I provide for this study will be treated confidentially
- I understand that in any report on the results of this research my identity will remain anonymous. This will be done by changing my name and disguising any details of my interview which may reveal my identity or the identity of people I speak about.
- I understand that disguised extracts from my interview may be quoted in:
    - In the dissertation of Yamini Srikanth
    - A final report to the forest department
    - In academic papers, policy papers or news articles
    - In an archive of the project as noted above
- I understand that signed consent forms, original audio recordings and transcripts, will be retained in Bengaluru India, accessible only by Yamini Srikanth and the research team, until three years post the completion of this interview.

**Signature of Research Participant** **Date**

______________________________

### Interview Consent Form (Tamil)

**நேர்காணல் ஒப்புதல் படிவம்**

**ஆராய்ச்சி திட்டத்தின் தலைப்பு** கப்பாஃபைகஸ் அல்வாரேசியின் சுற்றுச்சூழல் தொடர்புகள் மற்றும் தாக்கங்களை ஆய்வு செய்தல்

**புலனாய்வாளர்கள்:**
யாமினி ஸ்ரீகாந்த், உயிரியல் அறிவியலுக்கான தேசிய மையம், (, 9886901322)
Dr. Elrika D'Souza, Nature Conservation Foundation,
டாக்டர். சந்தீப் புள்ளா,
டாக்டர். நவீன் நம்பூத்ரி, தக்ஷின் அறக்கட்டளை.

**திட்டத்தின் தன்மை மற்றும் நோக்கம்:**

கப்பாஃபைகஸ் அல்வாரெஸி மன்னார் வளைகுடாவில் பயிரிடப்படும் ஒரு பொருளாதார முக்கியத்துவம் வாய்ந்த பாசி ஆகும். பவளப்பாறைகளில் அதன் எதிர்மறையான விளைவுகள் குறித்து கவலைகள் எழுந்துள்ளன. மன்னார் வளைகுடாவில் காணப்படும் தளங்களில் உள்ள ஒற்றுமைகள் மற்றும் வேறுபாடுகளைப் புரிந்துகொள்வதை நாங்கள் நோக்கமாகக் கொண்டுள்ளோம். நாங்கள் தாவர உண்ணிகளின் உண்ணும் முறைகள் பற்றியும் புரிந்து கொள்ள விரும்புகிறோம்.

**நேர்காணலின் தன்மை மற்றும் நோக்கம்:**

கப்பாஃபைகஸ் குருசடை மற்றும் ஷிங்கிள் தீவுகளில் முன்பு பரவியிருந்த அனைத்து இடங்களிலிருந்தும் மறைந்துவிட்டது. கடற்பாசி சேகரிப்பாளர்கள் பெரும்பாலும் விஞ்ஞானிகளை விட இந்த பகுதிகளுக்கு அடிக்கடி வருகை தருகின்றனர், மேலும் கடந்த சில ஆண்டுகளாக இந்த திட்டுகள் எவ்வாறு மாறிவிட்டன என்பதைப் பற்றிய ஆழமான புரிதல் உள்ளது. கப்பாஃபைகஸ் குருசடை மற்றும்
ஷிங்கிளில் இருந்து மறைந்தது மேலும் சாத்தியமான காரணங்களை மதிப்பிடவும் நாம் புரிந்து கொள்ள முயற்சி செய்கிறோம்,

நேர்காணல் முப்பது முதல் நாற்பத்தைந்து நிமிடங்கள் ஆகும். நீங்கள் அளிக்கும் பதிலினால் உங்களுக்கு எவ்வித பாதிப்பு ஏற்பட வாய்ப்பில்லை. எந்த நேரத்திலும் நேர்காணலை நிறுத்தவோ அல்லது ஆராய்ச்சியிலிருந்து விலகவோ உங்களுக்கு உரிமை உள்ளது.

மேலே உள்ள ஆராய்ச்சி திட்டத்தின் ஒரு பகுதியாக நேர்காணலுக்கு ஒப்புக்கொண்டதற்கு நன்றி. நேர்காணலுக்கு வருபவர்கள் நேர்காணலுக்கு வெளிப்படையாக ஒப்புக்கொள்வது மற்றும் அவர்களின் நேர்காணலில் உள்ள தகவல்கள் எவ்வாறு பயன்படுத்தப்படும் என்பது கல்விசார் ஆராய்ச்சிக்கான நெறிமுறை நடைமுறைகளுக்கு உட்பட்டது. உங்கள் ஈடுபாட்டின் நோக்கத்தை நீங்கள் புரிந்துகொள்வதையும், உங்கள் பங்கேற்புக்கான நிபந்தனைகளை நீங்கள் ஏற்றுக்கொள்கிறீர்கள் என்பதையும் உறுதிப்படுத்த, இந்த ஒப்புதல் படிவம் எங்களுக்கு அவசியம்.

**ஆராய்ச்சியில் பங்கேற்க ஒப்புதல்:**

- நான், ________________________________________ இந்த ஆராய்ச்சி ஆய்வில் பங்கேற்க தானாக முன்வந்து ஒப்புக்கொள்கிறேன்
- நான் இப்போது பங்கேற்க ஒப்புக்கொண்டாலும், எந்த நேரத்திலும் விலகலாம் அல்லது எந்த விதமான விளைவுகளும் இல்லாமல் எந்த கேள்விக்கும் பதிலளிக்க மறுக்கலாம் என்பதை நான் புரிந்துகொள்கிறேன்.
- நேர்காணலுக்குப் பிறகு இரண்டு வாரங்களுக்குள் யாமினி ஸ்ரீகாந்தைத் தொடர்புகொள்வதன் மூலம் எனது நேர்காணலிலிருந்து தரவைப் பயன்படுத்துவதற்கான அனுமதியை நான் திரும்பப் பெற முடியும் என்பதை நான் புரிந்துகொள்கிறேன், அப்படியானால் உள்ளடக்கம் நீக்கப்படும்.
- ஆய்வின் நோக்கம் மற்றும் தன்மையை எழுத்து மூலமாகவோ அல்லது வாய்மொழியாகவோ எனக்கு விளக்கியுள்ளன, மேலும் ஆய்வைப் பற்றி பதில் அளிக்க எனக்கு வாய்ப்பு கிடைத்தது.
- இந்த ஆராய்ச்சியில் பங்கேற்பதால் எனக்கு நேரடியாகப் பலன் கிடைக்காது என்பதைப் புரிந்துகொள்கிறேன்.
- எனது நேர்காணல் ஆடியோ பதிவு செய்யப்படுவதை ஒப்புக்கொள்கிறேன்.
- இந்த ஆய்வுக்காக நான் வழங்கும் அனைத்து தகவல்களும் ரகசியமாக கருதப்படும் என்பதை புரிந்துகொள்கிறேன்
- இந்த ஆராய்ச்சியின் முடிவுகள் குறித்த எந்த அறிக்கையிலும் எனது அடையாளம் பாதுகாப்பாக இருக்கும் என்பதை நான் புரிந்துகொள்கிறேன். இது எனது பெயரை மாற்றுவதன் மூலமும், எனது நேர்காணலின் எந்த விவரங்களையும் மறைப்பதன் மூலமும் எனது அடையாளத்தையோ அல்லது நான் பேசும் நபர்களின் அடையாளத்தையோ வெளிப்படுத்தாமல் செய்யப்படும்.
- எனது நேர்காணலில் இருந்து மறைக்கப்பட்ட பகுதிகள் மேற்கோள் காட்டப்படலாம் என்பதை நான் புரிந்துகொள்கிறேன்:
    - யாமினி ஸ்ரீகாந்தின் ஆய்வுக் கட்டுரையில்

- வனத்துறைக்கு இறுதி அறிக்கை
- கல்வித் தாள்கள், கொள்கைத் தாள்கள் அல்லது செய்திக் கட்டுரைகளில்
- மேலே குறிப்பிட்டபடி திட்டத்தின் காப்பகத்தில்

- கையொப்பமிடப்பட்ட ஒப்புதல் படிவங்கள், அசல் ஆடியோ பதிவுகள் மற்றும் மொழிபெயர்ப்பு,கள், இந்த நேர்காணல் முடிந்த மூன்று ஆண்டுகள் வரை, யாமினி ஸ்ரீகாந்த் மற்றும் ஆராய்ச்சிக் குழுவினரால் மட்டுமே அணுகக்கூடிய பெங்களூரு, இந்தியாவில் வைக்கப்படும் என்பதை நான் புரிந்துகொள்கிறேன்.

***ஆராய்ச்சி பங்கேற்பாளர் தேதியின் கையொப்பம்***

____________________________________________
